# The type I IFN-IL-27 axis promotes mRNA vaccine-induced CD8^+^ T cell responses

**DOI:** 10.1101/2025.01.16.633383

**Authors:** Anthony T. Phan, Emily Aunins, Elisa Cruz-Morales, Garima Dwivedi, Molly Bunkofske, Julia N. Eberhard, Daniel L. Aldridge, Hooda Said, Omar Banda, Ying Tam, David A. Christian, Robert H. Vonderheide, Ross M. Kedl, Drew Weissman, Mohamad-Gabriel Alameh, Christopher A. Hunter

## Abstract

The ability of lipid nanoparticle (LNP)-delivered mRNA vaccines to induce type I IFNs is critical to promote CD8^+^ T cell responses. The studies presented here indicate that immunization with nucleoside modified mRNA-LNP vaccines drives myeloid cell expression of the cytokine IL-27, which acts on antigen-specific CD8^+^ T cells to sustain T cell expansion. *In vitro* and *in vivo* studies revealed that type I IFN signaling is necessary for mRNA-LNP-induced IL-27 production, that immunization failed in IL-27 KO mice, and that immunization of IFNAR1-deficient mice with mRNA-LNP particles that also encode IL-27 mRNA restored antigen-specific CD8^+^ T cell responses. In addition, IL-27 mRNA-LNPs served as an adjuvant that improved cytolytic CD8^+^ T cell responses and the therapeutic efficacy of mRNA-LNPs to drive anti-pathogen and anti-tumor immunity. These studies highlight the central role of IL-27 in mRNA-LNP induced CD8^+^ T cell responses and the ability of this cytokine to augment the functionality of the CD8^+^ T cell response for prophylactic or therapeutic immunization.

## Introduction

Lipid nanoparticles (LNPs) formulated with N^1^-methyl-pseudouridine-modified mRNA (N1mΨ-mRNA) (mRNA-LNP) for vaccine antigen expression are potent inducers of protective immunity to SARS-CoV-2 infection and other pathogens^1–7^. The use of N1mΨ-mRNA is designed to reduce innate recognition of RNA and overt toxicity^8–10^, but mRNA-LNPs do stimulate the production of IL-6 and type I IFNs that contribute to the induction of humoral and cellular immunity^11,12^. The basis for mRNA-LNP induced CD8^+^ T cell responses is poorly understood. While type I IFNs are implicated in these events^11^, IL-6 and type I IFNs can promote or antagonize antigen-specific CD8^+^ T cells^13–21^. Here, we show mRNA-LNP vaccine-induced CD8^+^ T cell responses are not directly dependent on type I IFNs or IL-6 but rather that IL-27 acts directly on CD8^+^ T cells to support their expansion. Indeed, type I IFNs promote myeloid cell production of IL-27, which sustains the CD8^+^ T cell proliferative burst. Moreover, inclusion of IL-27 mRNA in mRNA-LNP formulations enhances CD8-mediated protective immunity to bacteria and tumors. These studies support the inclusion of cytokine mRNA-LNP adjuvants as a strategy to improve the development of prophylactic and therapeutic vaccines tailored to specific diseases.

## Results

### IL-27 is necessary for mRNA-LNP CD8^+^ T cell responses

Immunization with mRNA-LNP vaccines results in the rapid induction of inflammatory mediators that recruit immune cells and begins the processes required to generate a productive immune response^11,12,22^. In this context, the production of IL-6 and relatively low levels of type I IFNs contribute to the generation of protective antibody and antigen-specific CD8^+^ T cell responses respectively^11,12^. To assess what other inflammatory signals are induced by mRNA-LNP immunization, WT mice were immunized with PBS or a single 1 μg intramuscular (i.m.) dose of LNPs that contained N1mΨ-mRNA that encodes the model antigen ovalbumin (LNP-OVA). After 6 hrs, draining lymph nodes (dLN) were isolated and processed for multiplex cytokine analysis (Figure 1A). As expected^12^, IL-6 is highly induced (∼11x) with IL-27 (∼4.7x), IFN-γ (∼4.4x), and IL-12 (∼2x) the next three most-highly induced cytokines (Figure 1A). Previous studies found little role for endogenous IFN-γ and IL-12 in the ability of mRNA-LNP immunization to induce T cell responses^11,23,24^. IL-27, composed of the p28 and Ebi3 subunits, is a member of the IL-6/IL-12 family^25^, able to limit infection-induced effector CD4^+^ and CD8^+^ T cell responses and promote regulatory T cell functions^26–33^, yet IL-27 supports the induction of antigen-specific CD8^+^ T cell responses following subunit immunization^34,35^. Analysis of the dLN of immunized mice confirmed that mRNA-LNPs induced IL-27p28 as early as 4 hrs post-immunization with elevated levels through 48 hrs and a return to baseline by 72 hrs (Figure 1B). To identify the cellular source of IL-27, IL27p28GFP transgenic mice^35^, where *IL27p28* transcription results in expression of a green fluorescent protein (GFP), were immunized and the dLN analyzed at 24 and 48 hrs post-immunization. The numbers of immune cells that were GFP^+^ was elevated at 24 hrs and was further increased at 48 hrs post-immunization (Figure 1C-D). Analysis of the GFP^+^ cells identified inflammatory monocytes as the main source of IL-27, with the next most abundant cell types being type I (cDC1s), and type II dendritic cells (cDC2s) (Figure 1E-F). Vaccination did not increase systemic IL27p28 (Extended Data Fig. 1A) or drive GFP expression in contralateral lymph nodes of immunized mice (data not shown), suggesting that mRNA-LNP induced IL-27 is limited to the injection site draining lymph nodes at this dose of vaccine.

**Figure 1.**
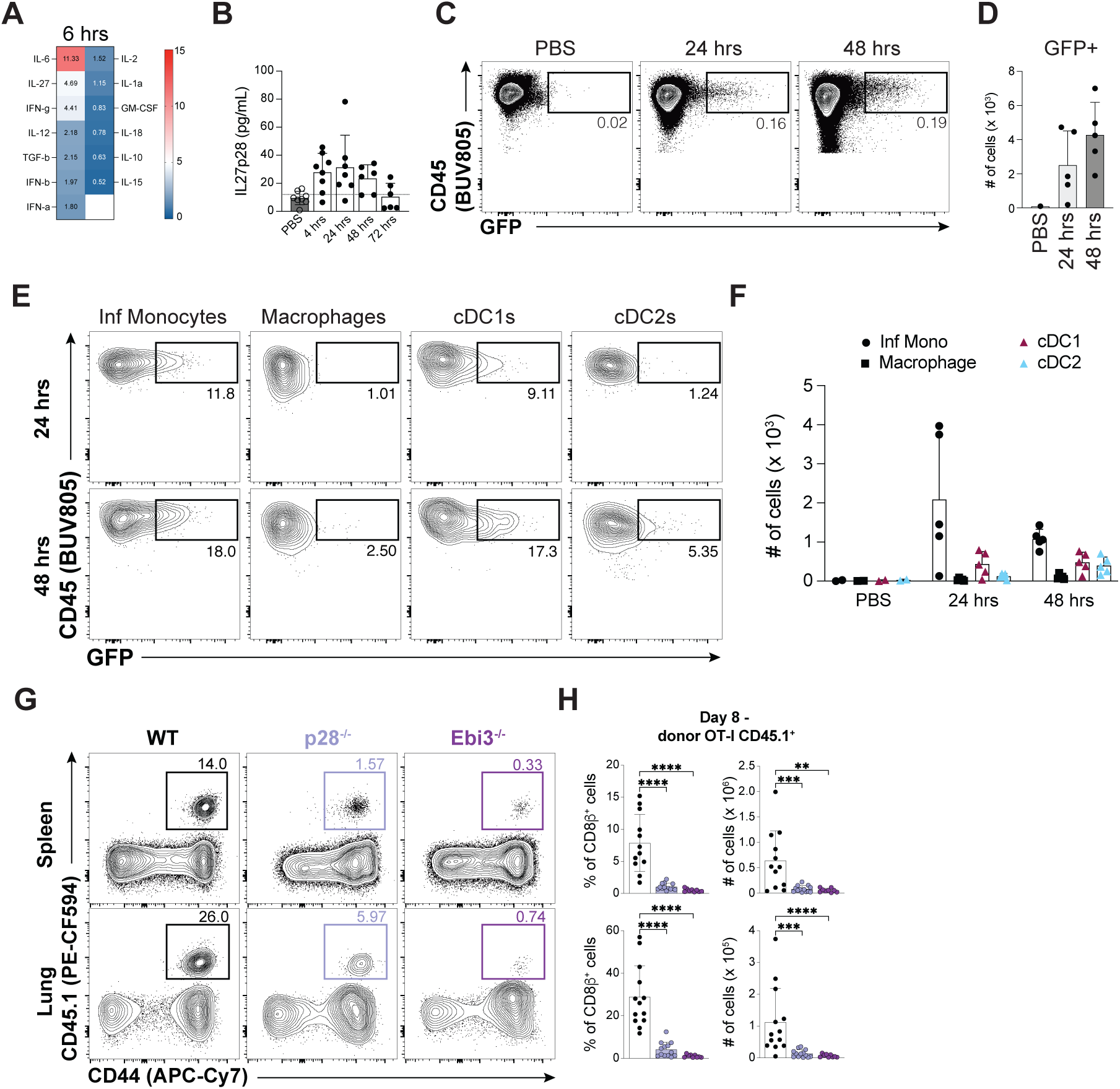
mRNA-LNP immunization promotes myeloid production of IL-27 necessary for CD8^+^ T cell responses. **A** Heat map of fold induction of indicated cytokines 6 hrs following mRNA-LNP immunization in the draining LNs of mice. Fold induction is the mean expression of mRNA-LNP immunized sample over PBS injected control samples. **B** IL-27p28 ELISA of draining lymph node lysates 4, 24, 48, and 72 hrs post-immunization with mRNA-LNPs. Dotted line is mean expression of IL27p28 in PBS injected mice. **C-D** Representative flow cytometry analysis of IL27p28-GFP expression of CD45^+^ cells in draining lymph nodes of mice at 24 (grey) and 48 (dark grey) hrs post-immunization **D** summarized absolute cell numbers. **E** Representative flow cytometry analysis of IL27p28-GFP expression by myeloid cells observed in draining lymph nodes 24 (top) and 48 (bottom) hrs post-immunization. **F** Absolute numbers of GFP^+^ myeloid cells in the draining lymph nodes of unimmunized mice, or mice 24 and 48 hrs post-immunization. **G-H** Flow cytometry analysis of adoptively transferred of WT OT-Is into WT (black), p28^-/-^ (light purple), or Ebi3^-/-^ (violet) host mice immunized i.m. with 1 μg LNP-OVA. **G** Representative flow cytometry analysis of donor OT-Is in spleen (top) and lungs (bottom) of indicated host mice 8 days post-immunization. **H** Summary proportion and number of OT-Is identified in spleen (top) and lungs (bottom) of indicated host mice 8 days post-immunization. WT, filled black circles, p28^-/-^, filled light purple circles, Ebi3^-/-^, filled violet circles. Multiplex cytokine array data are mean fold change of immunized mice over mice injected with PBS performed once (n = 6 mice immunized, n = 3 treated with PBS) **A**. Data are mean ± SD and representative (n = 3 – 5 mice per experimental group) **B-F** or summarized cumulative data (n = 3 – 4 mice per genotype) **H** from at least 2 independent experiments. ns p > 0.05, ** p < 0.01, *** p < 0.001, **** p < 0.001 for one-way ANOVA performed on **H**.

To assess the contribution of IL-27 to mRNA-LNP-induced CD8^+^ T cell responses, congenically distinct wildtype (WT) T cell receptor (TCR) transgenic OT-I CD8^+^ T cells (OT-Is) were adoptively transferred into WT, IL27p28^-/-^, or Ebi3^-/-^ mice, which are deficient in either IL-27 subunit, followed by immunization with LNP-OVA. At 8 days post-immunization, analysis of spleen and lungs of IL27p28^-/-^ and Ebi3^-/-^ host mice revealed the proportion and absolute numbers of OT-Is were significantly reduced in host mice that lacked IL-27 (Figure 1G-H). While IL-6 and IL-27 both utilize gp130 as part of their signaling receptor, and IL-6 can also promote CD8^+^ T cell activities^18,19,36^, when WT or IL-6^-/-^ hosts received OT-Is and were immunized with LNP-OVA there was no difference in the proportion and absolute numbers of responding OT-Is in the spleen and lungs (Extended Data Fig. 1B-C). Together, these data highlight that mRNA-LNP immunization induces IL-6 and IL-27 production in dLNs, but IL-27 has a distinct role in supporting subsequent CD8^+^ T cell responses.

### CD8^+^ T cell-intrinsic IL-27 signaling is necessary for mRNA-LNP-induced responses

Because the IL-27R is expressed by numerous innate and adaptive immune populations^37^, experiments were performed to determine whether the requirement for IL-27 for the CD8^+^ T cell responses described above was indirect or cell-intrinsic. Therefore, WT OT-Is or OT-Is that lack the IL-27 receptor alpha chain (IL27R^-/-^ OT-I) were individually transferred into congenically distinct WT mice followed by immunization with LNP-OVA (Figure 2A-C). Tracking the responding donor WT or IL27R^-/-^ OT-Is in peripheral blood over time revealed a defect in the response of IL-27R^-/-^ OT-Is that is apparent as early as day 6 (Extended Data Fig. 2A). At 8 days post-immunization, analysis of donor OT-Is confirmed the lack of IL27Ra expression by IL27R^-/-^ OT-I CD8^+^ T cells (Figure 2A) and revealed that IL27R^-/-^ OT-Is had a marked defect in expansion in spleen and lung (Figure 2B). Examination of phenotypic markers that delineate CD8^+^ T cell effector subsets (KLRG1, TCF1, CD127) showed that the proportion of TCF1^hi^CD127^hi^ (defining a vaccine-elicited subset^38^) or KLRG1^lo^CD127^hi^ (defining a subset of memory precursor effector cells^39^) cells within each donor population was similar irrespective of IL27R expression (Figure 2C and Extended Data Fig. 2B-C). Next, to determine if this defect is cell-intrinsic, a 1:1 mix of WT and IL27R^-/-^ OT-Is was transferred into WT hosts followed by immunization with LNP-OVA 1 day later. Tracking of responding donor CD8^+^ T cells in the peripheral blood of host mice confirmed that IL27R^-/-^ OT-Is exhibit a significant cell-intrinsic defect in their ability to proliferate following mRNA-LNP immunization whether assessed as a proportion of donor cells or host CD8^+^ T cells (Figure 2D-E). The defect in early expansion of IL27R^-/-^ OT-Is in comparison to WT OT-Is resulted in a significant reduction in the proportion and number of IL27R^-/-^ OT-Is observed in the spleen and lungs of host mice >30 days post-immunization (Extended Data Fig. 2D). Upon activation T cells rapidly alter their metabolic activity to accommodate the energetic requirements for a massive proliferative burst^40^. Previous studies highlighted that IL-27 signaling following subunit immunization is necessary for these metabolic adaptations and sustained proliferation^41^. Parallel studies establish that IL-27 promotes this via a hybrid transcriptional program unique to T cells responding to immunization composed of essential metabolic gene sets associated with expression of the transferrin receptor (CD71) and the large neutral amino acid transporter 1 (CD98)^38^. Indeed, when populations were assessed at day 4, a time point when it was already apparent that there were differences in expansion in the spleen (Figure 2F), WT OT-Is co-expressed high levels of CD71 and CD98 but this was significantly reduced in the absence of the IL27R (Figure 2G-H). These data support a model in which mRNA-LNP-induced IL-27 in the dLN directly signals to activated CD8^+^ T cells to sustain their proliferation.

**Figure 2.**
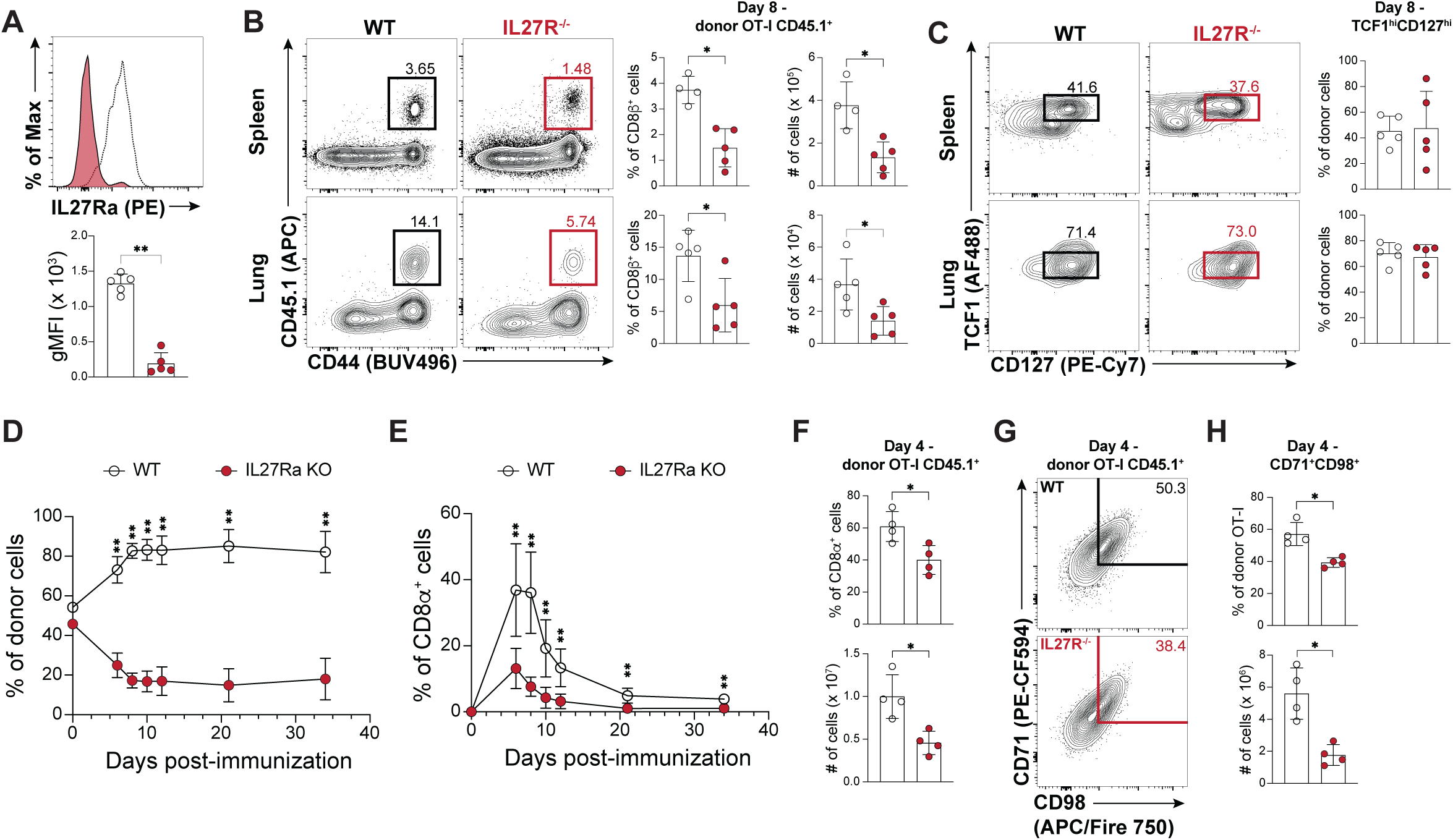
Direct IL-27 signaling is required for CD8^+^ T cell responses to mRNA-LNPs. **A-C** Flow cytometry analysis of WT host mice that received 10^3^ WT or IL27R^-/-^ OT-Is followed by i.m. immunization with 1 μg LNP-OVA. **A** Representative flow cytometry analysis of IL27Ra expression on WT or IL27R^-/-^ OT-Is found in the spleen of host mice day 8 post-immunization (open histogram = WT cells, filled histogram = IL27R^-/-^ cells). **B** Representative flow cytometry analysis of WT (open circles) and IL27R^-/-^ (red filled circles) OT-Is observed in spleen (top) and lungs (bottom) 8 days post-immunization. Summary proportion and number of WT and IL27R^-/-^OT-Is identified in spleen (top) and lungs (bottom) from wildtype host mice (right). **C** Flow cytometry analysis of TCF1 and CD127 expression of donor OT-Is in spleen (top) and lung (bottom) with summary graph of proportions of the gated subset (right). **D-E** Representative analysis of peripheral blood lymphocytes from WT mice that received a 1:1 mixed adoptive transfer of WT and IL27R^-/-^ OT-Is (10^4^ total cells) followed by i.m. immunization with 1 μg LNP-OVA. **D-E** Kinetic analysis of WT and IL27R^-/-^ OT-I CD8^+^ T cells in peripheral blood following mixed transfer as a proportion of donor cells **D** and as a proportion of host CD8^+^ T cells **E. F-H** Analysis of WT and IL-27R^-/-^ OT-I CD8^+^ T cells (10^4^ cells transferred) in spleen of wildtype host mice 4 days post-immunization. **F** Summarized proportion and absolute number of WT and IL27R^-/-^ OT-Is. **G-H** Representative analysis of CD71 and CD98 expression of WT (top) and IL27R^-/-^ (bottom) OT-Is in the spleen of host mice 4 days post-immunization. **G** Representative flow cytometry plots and **H** summary proportion (top) CD71^+^CD98^+^ donor OT-Is and number (bottom). Data are mean ± SD and representative from at least 2 independent experiments, n = 4 – 5 mice per genotype **A-C, F-H** and n = 5 – 10 for **D-E**. ns p > 0.05, * p < 0.05, ** p < 0.01, *** p < 0.001. Mann-Whitney non-parametric t-test used for **A-B, and F-H**. Multiple t-tests performed via Wilcoxon matched-pairs signed rank test corrected for FDR via Benjamini, Krieger, and Yekutieli method for **D-E.**

### Type I IFN supports LNP-induced CD8^+^ T cell responses via IL-27

Previous studies have highlighted a prominent role for type I IFNs in the CD8^+^ T cell response to intracellular infection and tumors^15,21,42^, but other reports indicated that direct type I IFN signaling was not necessary for adjuvant-elicited T cell responses^43,44^. These observations suggest that following mRNA-LNP immunization type I IFNs work in combination with IL-27, or the ability of type I IFNs to induce IL-27^45,46^ supports CD8^+^ T cell expansion. To distinguish between these possibilities, experiments were performed to determine whether mRNA-LNP induced type I IFNs contribute to the production of IL-27. When bone marrow-derived macrophages (BMDMs) from WT and interferon alpha receptor knockout (IFNAR^-/-^) mice were treated with mRNA-LNP particles, those from WT mice readily produced IL-27p28 whereas those from IFNAR^-/-^ mice produced significantly less (Figure 3A). Similarly, analysis of IL-27p28 production of immunized IFNAR^-/-^ mice revealed significantly impaired IL-27p28 production (Figure 3B). These data sets are consistent with a model in which mRNA-LNP induced type I IFN supports IL-27 production. However, analysis of IFN-α production in the dLNs following mRNA-LNP immunization of WT and IFNAR1^-/-^ mice revealed a significant defect in the ability of IFNAR1^-/-^ mice to produce IFN-α (Extended Data Fig. 3B). This observation concurs with reports that tonic type I IFN signaling supports expression of pattern recognition receptors^47–50^. Therefore, while IFNAR1^-/-^ mice immunized with mRNA-LNP have reduced vaccine-induced CD8^+^ T cell responses^11^, it remained unclear whether this required direct type I IFN signaling to CD8^+^ T cells.

**Figure 3.**
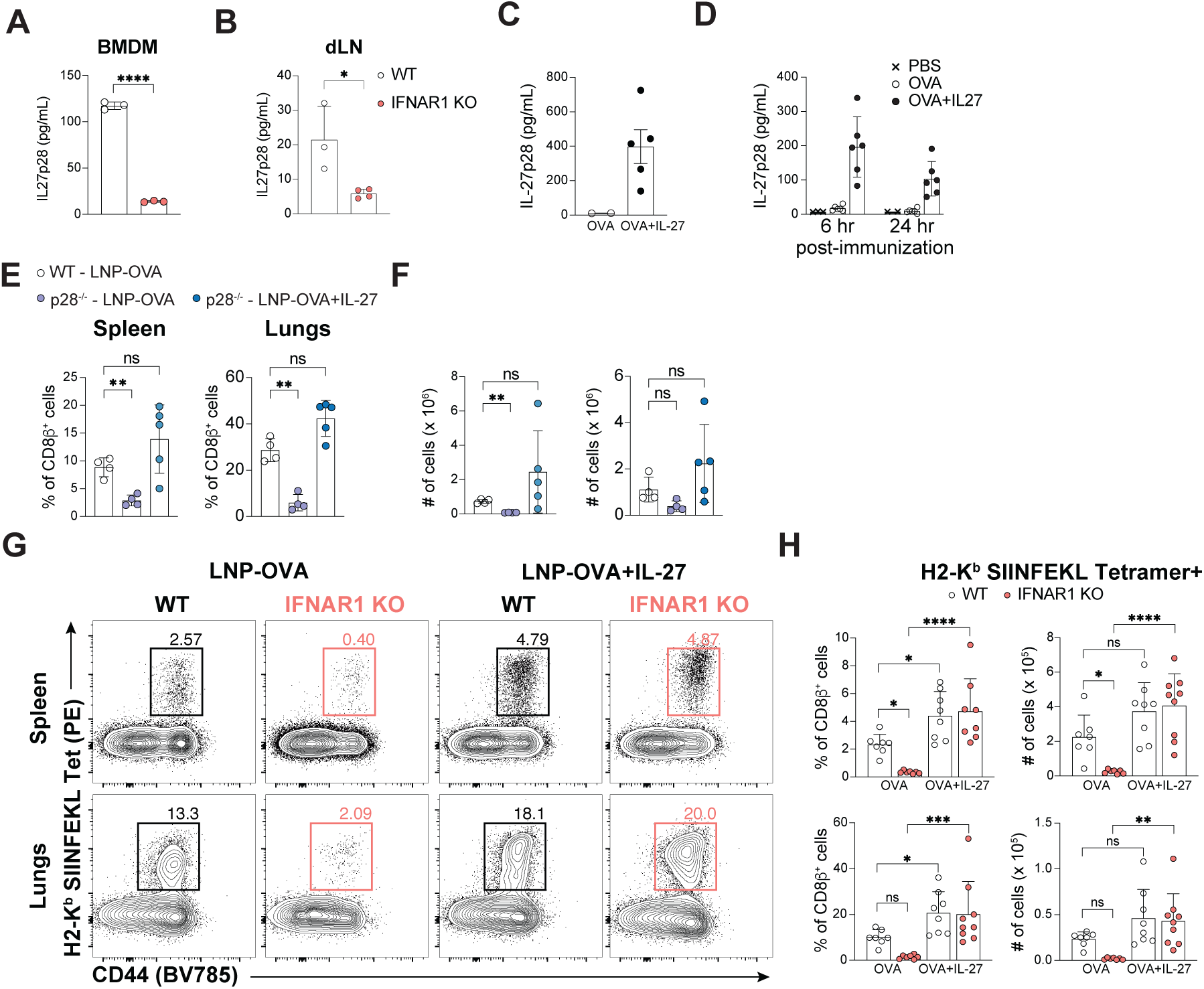
Type I IFN promotes CD8^+^ T cell responses to mRNA-LNP via IL-27. **A** IL27p28 ELISA of BMDM from WT and IFNAR1^-/-^ mice following overnight treatment with 1 μg LNP-OVA. **B** IL27p28 ELISA of dLN lysates from WT and IFNAR^-/-^ mice 4 hrs following immunization with LNP-OVA. **C-D** Measurement of IL-27p28 production by ELISA following treatment with LNP-IL-27 *in vitro* **C** and *in vivo* **D**. **C** IL-27p28 produced by BMDM treated with LNP-OVA or LNP-OVA+IL-27. **D** IL-27p28 expression in dLNs of mice immunized with LNP-OVA or LNP-OVA+IL-27 6 and 24 hrs post-immunization. **E-F** Summary graphs of flow cytometry analysis of WT OT-Is observed in spleens of WT or p28^-/-^ mice 8 days post-immunization with LNP-OVA or LNP-OVA+IL27 (WT immunized with LNP-OVA open circles, p28^-/-^ immunized with LNP-OVA filled light purple, p28^-/-^ immunized with LNP-OVA+LNP-IL27 filled light blue circles). **E** Proportion of donor OT-Is identified in spleen (left) and lungs (right) **F** Absolute numbers of donor OT-Is observed in spleen (left) and lungs (right)**. G-H** Flow cytometry analysis of antigen-specific CD8^+^ T cell response in WT and IFNAR^-/-^ mice 10 days post-immunization with 5 μg LNP-OVA or 5 μg LNP-OVA+IL-27. **G** Representative flow cytometry plots of activated H2-K^b^ SIINFEKL-specific CD8^+^ T cells in spleen (top) and lungs (bottom). H Summary proportion and absolute numbers of H-2K^b^ SIINFEKL-specific CD8^+^ T cells in spleen (top) and lungs (bottom) of WT and IFNAR1^-/-^ mice 10 days post-immunization with LNP-OVA or LNP-OVA+IL-27. WT, open circles, IFNAR1^-/-^, filled orange circles. Data are mean ± SD and representative **A-F** or summarized cumulative data **H** from at least 2 independent experiments. n = 3 – 5 mice per experimental group or genotype **B, D-H**. ns p > 0.5, * p < 0.5, ** p < 0.01, *** p < 0.001, and **** p < 0.0001. Unpaired t-test performed for **A-B**. Brown-Forsythe and Welch ANOVA tests followed by Dunnett’s T3 multiple comparison’s test performed for **E-H.**

To determine whether CD8^+^ T cell-intrinsic IFNAR1 signaling is necessary for responses to mRNA-LNP immunization, IFNAR1^-/-^ OT-I mice were generated (Extended Data Fig. 3C). Examination of responding WT or IFNAR1^-/-^ OT-I CD8^+^ T cells in the spleens of WT hosts 8 days post-immunization revealed that the IFNAR1^-/-^ OT-I CD8^+^ T cell response was indistinguishable from that of WT OT-Is (Extended Data Fig. 3D). To test whether the reduced IL-27 in the IFNAR1^-/-^ mice was responsible for limited CD8^+^ T cell responses, LNPs were formulated that contain N1mΨ-mRNA that encodes the IL27p28 and Ebi3 sub-units linked via a flexible glycine serine linker (Extended Data Fig. 3E)^51^. mRNA-LNPs that contained this construct drove the production of IL-27 in BMDM and *in vivo* (Figure 3C-D). Moreover, when WT OT-Is were adoptively transferred into WT or IL27p28^-/-^ mice that were then immunized with LNP-OVA, the inclusion of IL-27 mRNA rescued the OT-I responses in IL27p28^-/-^ mice to levels similar to WT mice immunized with LNP-OVA alone (Figure 3E-F). Next, WT and IFNAR1^-/-^ mice were immunized with LNP-OVA or LNP-OVA+IL-27, and H2-K^b^-SIINFEKL tetramer used to assess the polyclonal CD8^+^ T cell response. In WT mice, endogenous OVA-specific CD8^+^ T cell responses were improved when IL-27 mRNA was included in the vaccine formulation (Figure 3G-H). In line with previous studies, IFNAR1^-/-^ mice exhibited a significant defect in their OVA-specific CD8^+^ T cell response but this defect was rescued by the inclusion of the IL-27 mRNA (Figure 3G-H). These data sets indicate that type I IFN signaling promotes the production of IL-27 which supports the expansion of the CD8^+^ T cell response and that provision of IL-27 is sufficient to override the requirement for type I IFNs (Extended Data Fig. 3F).

### Co-delivery of IL-27 mRNA enhances CD8^+^ T cell responses to LNP-mRNA vaccination

The data above indicate the necessity for IL-27 to sustain CD8^+^ T cell expansion in contrast to other cytokines such as IL-12 that promote expansion and differentiation of effector^21^ CD8^+^ T cells yet may limit memory differentiation^52^. To evaluate the impact of LNP-OVA+IL-27 on the CD8^+^ T cell effector and memory responses OT-Is were transferred into WT hosts prior to immunization with a 1 μg dose of LNP-OVA+IL-27 or LNP-OVA that was mixed with an irrelevant mRNA (irrmRNA), to ensure equivalent mRNA and LNP dose, and T cell responses assessed at 8 and 30 days post-immunization (Figure 4A-F). Analysis of the OT-I response in spleen and lung at day 8 revealed that inclusion of IL-27 resulted in an increased population of vaccine-induced OT-Is (Figure 4A) that retained high expression of CD127 (IL-7Ra), lacked KLRG1 expression, and did not alter tissue distribution within the spleen or lungs (Figure 4B-C). By 30 dpi, this T cell response had contracted and the number of OT-Is trended higher in mice treated with IL-27 (Figure 4D-E). This was independent of the LNP formulation as similar results were observed when utilizing LNPs formulated with an alternative ionizable lipid SM-102 when tracking vaccine-elicited OT-Is via peripheral blood (Extended Data Fig. 4A-B).

**Figure 4.**
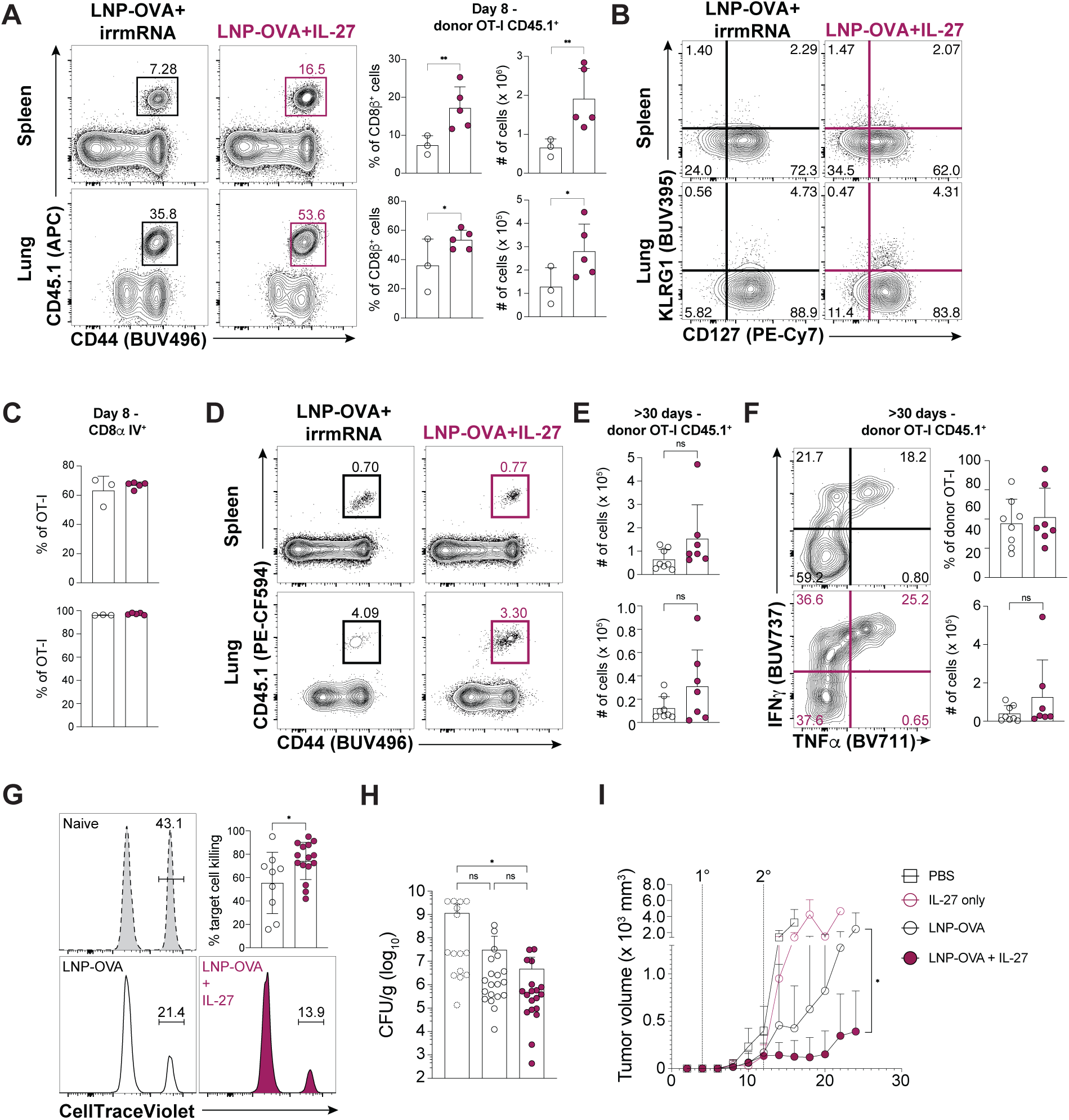
Addition of IL-27 mRNA during LNP-mRNA dosing enhances CD8^+^ T cell responses. **A-C** Representative flow cytometry analysis of WT OT-I CD8^+^ T cells observed in spleen and lungs of wildtype host mice immunized with LNP-OVA+irrmRNA or LNP-OVA+IL-27 8 days post-immunization. **A** Proportion and absolute numbers of WT OT-I CD8^+^ T cells identified in spleen (top) and lungs (bottom). Summary graphs of proportion and absolute numbers (right). **B** KLRG1 and CD127 cell surface phenotype of donor OT-Is observed in spleen and lungs. **C** Proportion of donor OT-Is labeled with intravenous injected CD8α antibody 8 days post-immunization in spleen (top) and lungs (bottom). **D-F** Representative cytometry analysis of donor OT-Is identified >30 days post-immunization in spleen (top) and lungs (bottom) of wildtype mice immunized with 1 μg dose of LNP-OVA+irrmRNA or LNP-OVA+IL-27. **E** Summary graphs of absolute number of donor OT-Is observed >30 days post-immunization. **F** Representative IFNγ and TNFα production of donor OT-Is in the spleen of mice immunized with LNP-OVA+irrmRNA (top) or LNP-OVA+IL-27 (bottom) following 4 hrs of restimulation with SIINFEKL peptide. Summary proportion and absolute numbers of IFNγ^+^ cells (right). **G** Histograms of CellTraceViolet labeled target cells in naive (dashed line grey filled) mice, LNP-OVA immunized (open), or LNP-OVA+IL-27 (maroon filled) immunized >30 days prior. Summarized proportion of target cell killing (right). **H** CFU/g of Listeria monocytogenes Ovalbumin 66 hrs post challenge of naive (dashed open circle), 1 μg LNP-OVA (open), 1 ug LNP-OVA+IL-27 (maroon filled). **I** Tumor volume on right hind flank of mice that received initial subcutaneous inoculation of 2 x 10^5^ B16F0-OVA and treated with PBS (open square), LNP-IL-27 (open maroon circle), LNP-OVA (open circle), or LNP-OVA+IL-27 (filled maroon circle) at 4 and 12 days post tumor inoculation over time. Data are mean ± SD and representative **A-C,I** or summarized cumulative **E-H** from at least 2 independent experiments. n = 3 – 5 mice per experimental group **A-F**, n = 5 – 10 mice per experimental group for **G-H**. n = 5 mice IL-27 only, n = 10 for PBS, LNP-OVA, and LNP-OVA+IL-27 treated for **I**. ns p > 0.05, * p < 0.05, ** p < 0.01, *** p < 0.001. Mann-Whitney non-parametric t-test performed for **A-F, H**. Unpaired student’s t-test performed for **G.** Mixed-effects analysis with Tukey’s test for multiple comparisons for **I**.

Importantly, though the increased expansion at day 8 (Figure 4A) did not impair the generation of longer-lived antigen-specific CD8^+^ T cells in mice that received LNP-OVA+IL-27. Antigen-specific CD8^+^ T cells responding to LNP-OVA+IL-27 immunization maintained their effector function at >30 dpi. Moreover, following peptide restimulation (>30 dpi) the ability to produce IFN-γ and TNFα was assessed by intracellular staining. OT-Is from LNP-OVA+IL-27 treated mice were still able to effectively produce these cytokines (Figure 4F). In addition, an *in vivo* CTL assay revealed that mice immunized with a single dose of LNP-OVA were able to kill ∼50% of peptide-pulsed target cells, but when IL-27 was included this resulted in significantly improved killing of target cells despite modest increases in SIINFEKL-tetramer^+^ CD8^+^ T cells (Figure 4E and Extended Data Fig. 4C). To determine if the improved killing of target cells translated to improved secondary responses to infection, mice immunized ≥30 days prior were challenged intravenously with *Listeria monocytogenes* that express ovalbumin (Lm-OVA) and >60 hrs later bacterial burden was assessed (Figure 4H). As expected, naive mice had a high bacterial burden, and immunization with LNP-OVA resulted in a reduced CFU/g that was further reduced with the inclusion of IL-27 mRNA (Figure 4H). These data demonstrate IL-27 mRNA improves the protective capacity of antigen-specific cells following a single immunization (Figure 4A-H).

Finally, we assessed whether LNP-OVA+IL-27 could be useful as a therapeutic vaccine in cancer. B16F0-OVA melanoma tumor cells were implanted subcutaneously in the flanks of WT mice and then randomly assigned into treatment groups. Tumor implanted mice were then treated with an i.m. dose of PBS, LNPs formulated with IL-27 mRNA only (LNP-IL-27), LNP-OVA, or LNP-OVA+IL-27 at day 4 and day 12 post-tumor inoculation (Figure 4I) and tumor growth assessed. In mice that only received PBS or LNP-IL-27 tumors grew rapidly while those that received LNP-OVA, tumor growth was slowed initially but broke through by day 16 post-inoculation and then showed progression. In contrast, mice that received LNP-OVA+IL-27 showed significantly slowed tumor growth throughout the course of the study (Figure 4I). The ability of an IL-27 mRNA adjuvant to improve tumor control highlights the promise of utilizing cytokine encoding mRNAs in LNP formulations and that inclusion of IL-27 mRNA has potential to improve cellular immunity in prophylactic and therapeutic vaccines for infectious diseases and oncologic applications.

## Discussion

Despite their success and widespread use, an understanding of the cellular and molecular mechanisms through which mRNA-LNP vaccines potentiate protective responses remains unclear. Here we identify that vaccination with mRNA-LNPs leads to the local induction of type I IFNs that in turn promote the production of IL-27 that serves an essential role in promoting CD8^+^ T cell responses. In addition, the inclusion of IL-27 mRNA in vaccine formulations is sufficient to override the requirement for type I IFN to induce CD8^+^ T cell responses. These data, along with those in the concurrent study^38^, support a role for IL-27 signaling in maintenance of an early transcriptional program that is necessary for maximal CD8^+^ T cell proliferation following vaccination^34,41^. These data sets are consistent with reported roles for IL-27 to sustain CD8^+^ T cell function in persistent viral infections^53^, as well as with a report that patients that lack IL-27R have reduced ability to generate Epstein-Barr virus-specific cytotoxic CD8^+^ T cells^54^. Likewise, there are reports of patients with auto-antibodies to IL-27^54^ or type I IFNs^55–57^ that are more susceptible to viral infections. Specifically, vaccinated patients with high levels of auto-antibodies to type I IFNs can generate relatively normal levels of SARS-CoV-2-specific neutralizing antibodies, but exhibit significantly higher risk of severe breakthrough SARS-CoV-2 infections^57^. This observation is reminiscent of the ability of type I IFNs in mice to promote CD8^+^ T cell responses to SARS-CoV-2 immunization^11^ and suggests these patients could fit a model in which their reduced type I IFN signaling impaired the generation of CD8^+^ T cells capable of providing protection from SARS-CoV-2.

While mRNA vaccine efficacy hinges on immune pathways that operate during infection (i.e. IL-6, type I IFN signaling, IL-27) – the specifics of when and how they operate are critical for the breadth and protective capacity of the humoral and cellular immune responses. For example, IL-27 can be profoundly suppressive of CD8^+^ T cell responses during certain infections^27,28,58,59^, but in the data sets presented here and in the context of subunit vaccination strategies it supports CD8^+^ T cell expansion^34,41^. One implication of these and other studies is that the outcome of mRNA-LNP vaccination hinges on a complex network of inflammatory signals that promote humoral or cellular immunity which can be modified by the inclusion of mRNA-encoded adjuvants. Thus, the ability to bypass the requirement for type I IFN by inclusion of mRNA encoding a cytokine highlights the potential to engineer or tailor the CD8^+^ T cell response to generate improved cytotoxic responses to infection and malignancy, thereby broadening the potential therapeutic use of mRNA-LNPs^60,61^. Moreover, the modular mRNA-LNP platform allows for creative application and further iteration that includes the use of mRNA for pattern recognition receptors as well as other cytokines ^23,24,62,63^. The ability to incorporate additional immunomodulatory molecules allows flexibility for the design of novel vaccines and therapies.

## Materials and Methods

### Mice

IL27Ra deficient (IL27Ra^-/-^, JAX stock #018078)^64^, IFNAR1 deficient (IFNAR1^-/-^, JAX stock #028288), OT-I TCR transgenic mice (JAX stock #003831)^65^, B6.SJL-Ptprca Pepcb/BoyJ (CD45.1, JAX stock #002014), C57BL/6J-Ptprcem6Lutzy/J (JAXboy, JAX stock #033076), and wildtype C57BL6/J (B6, JAX stock #000664) were purchased from Jackson Laboratory and housed in the University of Pennsylvania School of Veterinary Medicine vivarium for at least 1 week prior to studies being performed. IL27Ra^-/-^ and OT-I mice crossed to CD45.1 mice were crossed to generate IL27Ra^-/-^ OT-I CD45.1 or CD45.1.2 mice. IFNAR1^-/-^ mice were crossed to OT-I mice to generate IFNAR1^-/-^ OT-I mice. OT-I mice were crossed to CD45.1 or JAXboy mice to generate OT-I CD45.1.2 or CD45.1 mice. IL27p28^-/-^ and Ebi3^-/-^ mice on C57BL/6J background were generated by Lexicon Genetics. For all animal studies male and female mice 6-12 weeks of age were used. All mice were housed in a specific-pathogen free environment at the University of Pennsylvania School of Veterinary Medicine in accordance with federal guidelines and with approval of the Institutional Animal Care and Use Committee.

### Immunization and mRNA-LNPs

Codon-optimized sequences for Ovalbumin (OVA) and glycine serine linked murine Ebi3 with murine IL27p28 (IL-27) were gene synthesized and cloned into an mRNA production plasmid and mRNA was produced as previously described^66^. mRNA was purified by cellulose purification, analyzed by agarose gel electrophoresis and stored frozen at -20 C. Cellulose-purified N1mΨ-containing RNAs were encapsulated in LNPs using a self-assembly process previously described^67^. The LNP formulation used in this study is proprietary to Acuitas Therapeutics; the proprietary lipid and LNP composition are described in US patent US10,221,127. SM-102 LNPs were in house formulated using the Ignite Nanoassembler microfluid device. An ethanolic mixture of SM-102, DSPC, Cholesterol and PEG-Lipid at 50:10:38.5:1.5 % mol/mol was prepared, and rapidly mixed with an aqueous solution of mRNA, in line diluted, dialyzed, and concentrated to a final concentration of 1mg/mL. All LNPs were around 80 nm in diameter, had an encapsulation efficiency > 95%, and displayed a zeta potential of around -2 to -5 mV. All mRNA-LNPs were stored at -80°C until use. All mice were immunized intramuscularly in the right hind leg with 1 μg of indicated mRNA-LNP diluted with PBS to a volume of 50 μl, unless otherwise indicated. When mRNA-LNPs are co-administered, dosage was a 1:1 mix of mass:mass of indicated mRNA-LNP formulations.

### T cell transfers

For T cell transfer studies splenocytes from OT-I TCR transgenic mice of necessary genotypes were enriched by magnetic activated cell sorting (MACS) using the CD8a^+^ T Cell Isolation Kit (Miltenyi Biotec, 130-104-075), or EZ Sep CD8^+^ T cell enrichment kit (StemCell Technologies, 19853) following manufacturer instructions. Enriched cells were counted and then indicated numbers of cells (10^3^ – 10^4^) were transferred via retro-orbital injection one day prior to immunization with mRNA-LNPs. Enriched cells were used for mixed adoptive transfers. Pre-transfer mixes of OT-I CD8^+^ T cells were confirmed via flow cytometry prior to injection to record starting ratio of transferred cells.

### Tissue harvest and preparation

#### IV labeling of circulating lymphocytes

Circulating lymphocytes were labeled by intravenous injection of 3 μg of indicated antibodies (specific for CD8a or CD45) retro-orbitally 3 minutes prior to euthanasia of the animal as previously described^68^.

#### Secondary lymphoid tissues

Single cell suspensions from draining lymph nodes (inguinal and popliteal ipsilateral to injected muscle) and spleen were isolated by processing tissues over a 70 μm cell strainer (Fisher Scientific, 22-363-548), washing with RPMI (Corning, 10-040-CM) supplemented with 5% Fetal Bovine Serum (RP-5), followed by red blood cell lysis with 1X ACK lysis buffer for 3 min at room temperature, washing cells once more, then resuspending in fresh RP-5.

#### Lungs

Single cell suspensions from the lung were obtained by digesting chopped lung tissue in 4 mL of RPMI supplemented with 2.5% FBS, Type IV collagenase (400 U/mL) and DNase I (0.33 μg/mL) for 30-40 min in a shaking incubator at 37°C. Digested tissue was then processed through a 70 μm filter and washed with RP-5. Red blood cells were lysed as above and then cells were washed and resuspended in fresh RP-5.

All processed single cell suspensions were counted on a Guava HTS with ViaCount reagent for absolute numbers of viable cells.

#### Peripheral blood

150 μl of blood was collected at indicated time points following immunization from mice via submandibular bleeding into lithium heparin treated tubes (BD #365965). Red blood cells were lysed with 1X ACK lysis buffer at room temperature. Samples were resuspended in 200 μl of FACS staining buffer and a sample was counted on the Guava HTS with ViaCount reagent while the remainder was stained for flow cytometry analysis.

#### Draining Lymph Node Lysates

All samples were kept on ice throughout the process. Draining lymph nodes were harvested into ZR BashingBead (2 mm) Lysis Tubes (Zymogen, S6003-50) with 300-600 μl of EZLys Tissue Protein Extraction Reagent (BioVision, 8002-500) supplemented with 1X HALT Protease and Phosphotase Inhibitor Cocktail (Thermofisher Scientific, 78440).

### Flow cytometry

After processing, equivalent numbers of cells were plated and stained for viability using Ghost Dye Violet 510 (Tonbo Biosciences, 13-0870-T100) or Ghost Dye Red 780 (Tonbo Biosciences, 13-0865-T100) for 20 min at 4°C. Cells were washed with FACS staining buffer (FSB, 1X PBS supplemented with 0.2% bovine serum albumin (Gemini, 700-100P), and 1 mM EDTA (Gibco, 15575-038)) then incubated in Fc block (FSB supplemented with 1 μg/mL anti-mouse CD16/CD32 (Bio X Cell, BE0307), 3 μg/mL normal Mouse IgG (Thermofisher Scientific, 10400C), 3 μg/mL normal Rat IgG (Thermofisher Scientific, 10700) for 10 min at 4°C. Tetramer-specific CD8^+^ T cells were identified by staining in 50 μl of FSB containing H2-K^b^ SIINFEKL tetramers provided by the NIH tetramer core for 35-45 min at 4°C. Cells were then surface-stained in 50 μl at 4°C for 30 min and washed before acquisition if intracellular staining was not performed. For detection of intracellular transcription factors cells were fixed and stained with the eBioscience Foxp3 Transcription Factor Staining Kit (ebioscience, 00-5523-00) according to manufacturer’s instructions. For intracellular cytokine staining, cells were washed with FSB following surface staining and then fixed and stained with the BD Cytofix/Cytoperm Kit (BD Biosciences, 554714) according to manufacturer’s instructions. All flow cytometry data acquisition was performed on BD Symphony A3 or A5 instruments maintained by Penn Cytomics and Cell Sorting Shared Resource Laboratory. Data analysis was performed in FlowJo v10.10 (BD) with data cleanup performed with FlowAI plugin to eliminate poor quality events^69^.

### Quantification of mRNA-LNP induced cytokines

In brief supernatants from *in vitro* cultured bone marrow derived macrophages (BMDMs) of indicated genotypes were collected or draining lymph nodes, inguinal and popliteal, were harvested from immunized mice at indicated time points and lysed for tissue lysates as detailed above. Supernatants or tissue lysates were assayed by ELISA for IL-27p28 and IFN-α (R&D Biosystems Cat# M2728 and MFNAS0 respectively) according to manufacturer’s instructions. Multiplex cytokine assay for screening of cytokines induced by mRNA-LNP immunization *in vivo* was performed with a custom LegendPlex panel (Biolegend Cat# 900001953) according to manufacturer’s instructions. Data from LegendPlex screening displayed as mean fold change of each immunized mouse over mean cytokine expression from PBS-treated draining LN lysates for each cytokine examined. All samples were aliquoted and frozen down at -80°C until thawed for assay.

### In vivo Cytotoxic Lymphocyte (CTL) Assay

Congenically distinct splenocytes were either pulsed with SIINFEKL peptide for 1 hour at 37°C or left unpulsed and labeled with high (1:2000 dilution) or low concentrations (1:40,000 dilution) of CellTraceViolet (Invitrogen, C34557) respectively. A 1:1 mix of cells was then transferred intravenously via lateral tail-vein injection into naive or mice vaccinated at least 30 days prior. 16 hours post-transfer mice were euthanized and spleens harvested and processed to assess target cell killing via the ratio of pulsed and unpulsed cells recovered from each spleen.

### Tumor Cell Lines

B16F0-OVA cell line was provided by Dr. Mohamad-Gabriel Alameh (University of Pennsylvania). Tumor cells were grown in DMEM supplemented with 10% FBS, 10 mM HEPES, 1 mM sodium pyruvate, 1X GlutaMAX, 1x MEM Non-essential amino acids, 55 μM β-mercaptoethanol, 1X Penicillin/Streptomycin and G418 (400 μg/mL).

### Melanoma tumor model and therapeutic immunization

The right flanks of study mice were shaved at least 2 days prior to tumor implantation to allow easier measurement of subcutaneous tumor growth. 2 x 10^5^ B16F0-OVA cells were injected subcutaneously (s.c.) into the flank in 200 μl of sterile Hank’s Balanced Salt Solution. Therapeutic immunizations (10 μg total vaccine dose of indicated treatments or 50 μl PBS) were administered to groups of randomly assigned mice i.m. on day 4 and day 12 post-tumor cell injection. Growth of tumors was monitored every 2-3 days for at least 24 days or until humane endpoint (tumor volume of 4000 mm^2^ or if ulceration is observed). Tumor volumes were measured by using vernier caliper and calculated with the following equation: V = (4 x 3.14 x A x B^2^)/3, where V = volume (mm^3^), A = the largest diameter of the tumor (mm), and B = the smallest diameter of the tumor (mm).

## Statistical analysis

Indicated statistical analyses were performed using Prism software (GraphPad Software LLC). Tests of normality and lognormality were run on all statistically analyzed data sets to aid in the selection of statistical analyses used. Significance was calculated with appropriate statistical tests and post-tests as listed in figure legends.

## Data availability

Data supporting the findings of this study are available within the article and the main figures or it’s extended data. Raw flow cytometry data presented in the figures and used to generate the plots are available upon request from the corresponding author C.A.H..

## Acknowledgements

Funding of this work provided by the US NIH (U19-AI142596, U19-AI142596, UM1-AI144371, 75N93019C000550, and P01AI158571) for M-G.A, G.D., and D.W., US NIH (UO1 AI160664) for R.M.K., US NIH (UO1 AI160664, AI157247) for C.A.H., the Commonwealth of Pennsylvania through the Institute of Infectious and Zoonotic diseases for C.A.H., the Basser Interception Center for C.A.H., D.W., and R.H.V., the Parker Institute for Cancer Immunotherapy for R.H.V., the Emerson Collective for A.T.P. and C.A.H., US NIH Cancer T32 (CA009140) for M.B., US NIH Ruth L. Kirschstein Predoctoral Fellowship (F31AI161962) for D.L.A.. We thank the Penn Institute for RNA Innovation for funding. We thank the Penn Cytomics & Cell Sorting Shared Resource Laboratory as flow cytometry data in this manuscript was generated on their instruments and is partially supported by the Abramson Cancer Center NCI Grant (P30 016520), research identifier number RRid:SCR_022376. We thank C. Lombana and the Scott lab for providing recombinant Listeria-OVA. We thank R. Pardy, Z. Lanzar, and Hunter lab members for critical discussions and review of the manuscript; and Q. Fang for technical assistance.

## Author Contributions

Conceptualization: A.T.P., and C.A.H.

<u>Methodology:</u> A.T.P., E.A., E.C-M., H.S., O.B., G.D., M.B., J.N.E., R.M.K., and M-G.A.

<u>Analysis:</u> A.T.P., E.A., E.C-M., G.D., and M-G.A.

<u>Investigation:</u> A.T.P., E.A., E.C-M., G.D., M.B., J.N.E., D.L.A., D.A.C., and M-G.A.

<u>Resources:</u> Y.T., R.M.K., M-G.A., D.W., and C.A.H.

<u>Writing - Original Draft:</u> A.T.P and C.A.H.

<u>Writing – Review and Editing:</u> All Authors

<u>Supervision:</u> M-G.A., and C.A.H.

<u>Funding Acquisition:</u> R.M.K., D.W., M-G.A., R.H.V., and C.A.H.

## Competing Interests Statement

In accordance with the University of Pennsylvania and Children’s Hospital of Philadelphia policies and procedures and our ethical obligations as researchers, we report that A.T.P, E.A., M-G.A., D.W., and C.A.H. have filed a provisional patent application that describes the use of nucleoside-modified mRNA encoding cytokines as adjuvants. D.W. is named on patents that describe the use of nucleoside-modified mRNA as a platform to deliver therapeutic proteins and vaccines. D.W. and M-G.A. are named on patents describing the use of lipid nanoparticles, and lipid compositions for nucleic acid delivery and vaccination. We have disclosed those interests fully to the University of Pennsylvania and the Children Hospital of Philadelphia, and we have in place an approved plan for managing any potential conflicts arising from licensing of our patents. Y.T. is an employee of Acuitas Therapeutics and is named on patents describing the use of lipid nanoparticles for nucleic acid delivery. M-G.A. serves as a scientific adviser for AfriGen Biologics and has an ownership stake in RNA Technologies. D.W. serves as a scientific advisor for Arcturus Therapeutics, Cabaletta Bio, and Versatope Therapeutics, and has ownership stakes in Capstan Therapeutics, Orbital Therapeutics, Zipcode Bio, and RNA Technologies. D.W. receives royalties from CellScript and Capstan Therapeutics. R.H.V. has received consulting fees from BMS and Grey Wolf Therapeutics, research funding from Revolution Medicines, is an inventor on patents relating to cancer cellular immunotherapy, cancer vaccines, and KRAS immune epitopes, and receives royalties from Children’s Hospital Boston for a licensed research-only monoclonal antibody. The remaining authors declare no competing interests.

## Extended Data legends

**Extended Data Figure 1.**
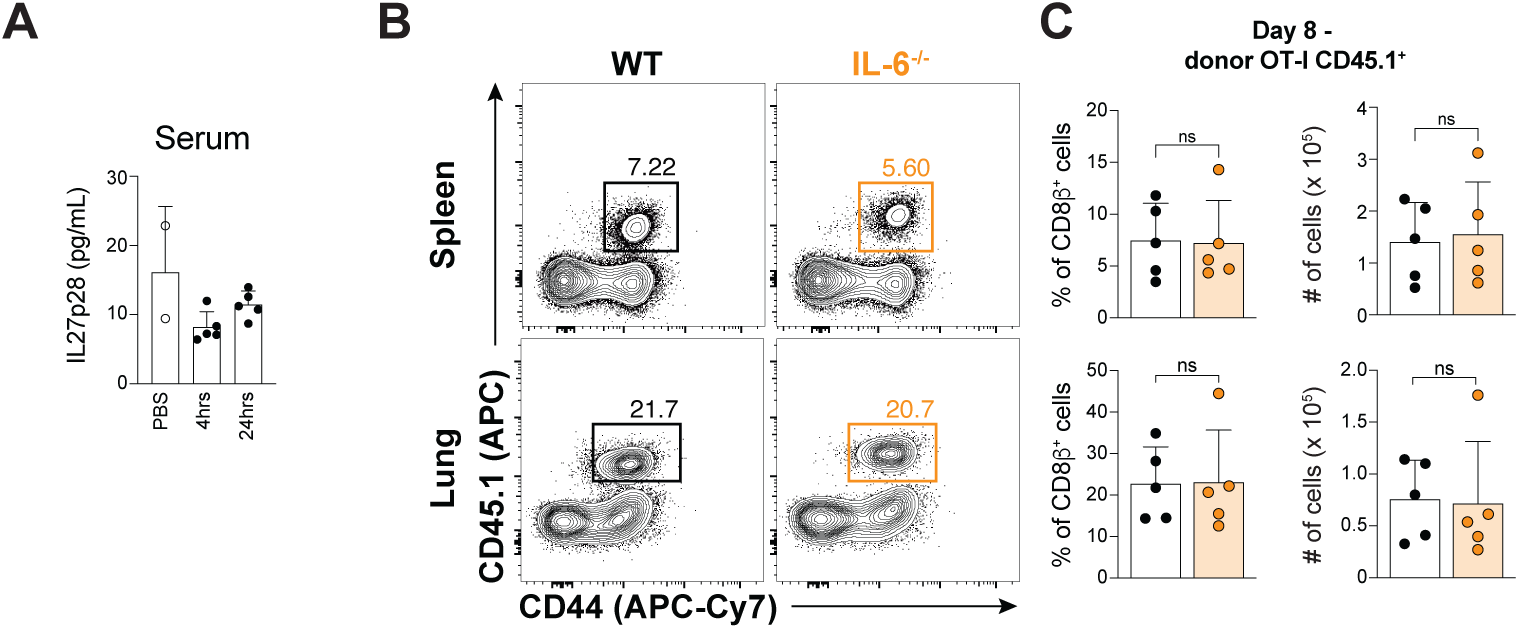
Supporting data that local mRNA-LNP induced IL-27 is uniquely required for CD8^+^ T cell responses. **A** IL-27p28 ELISA performed on serum samples from B6 mice immunized with 1 μg LNP-OVA 4 hrs and 24 hrs post-immunization. **B-C** Flow cytometry analysis of adoptively transferred OT-Is observed in spleen and lungs of WT and IL-6^-/-^ host mice 8 days post-immunization. **B** Representative flow cytometry plots of donor OT-Is identified in spleen (top) and lungs (bottom) of WT (black) and IL-6^-/-^ (orange) host mice. **C** Summary graphs of proportion and absolute number of observed donor OT-Is. Representative data from studies performed 2 times with n = 5 mice for indicated experimental groups. Data are mean ± SD. ns, p value > 0.05 following Mann-Whitney non-parametric t-test.

**Extended Data Figure 2.**
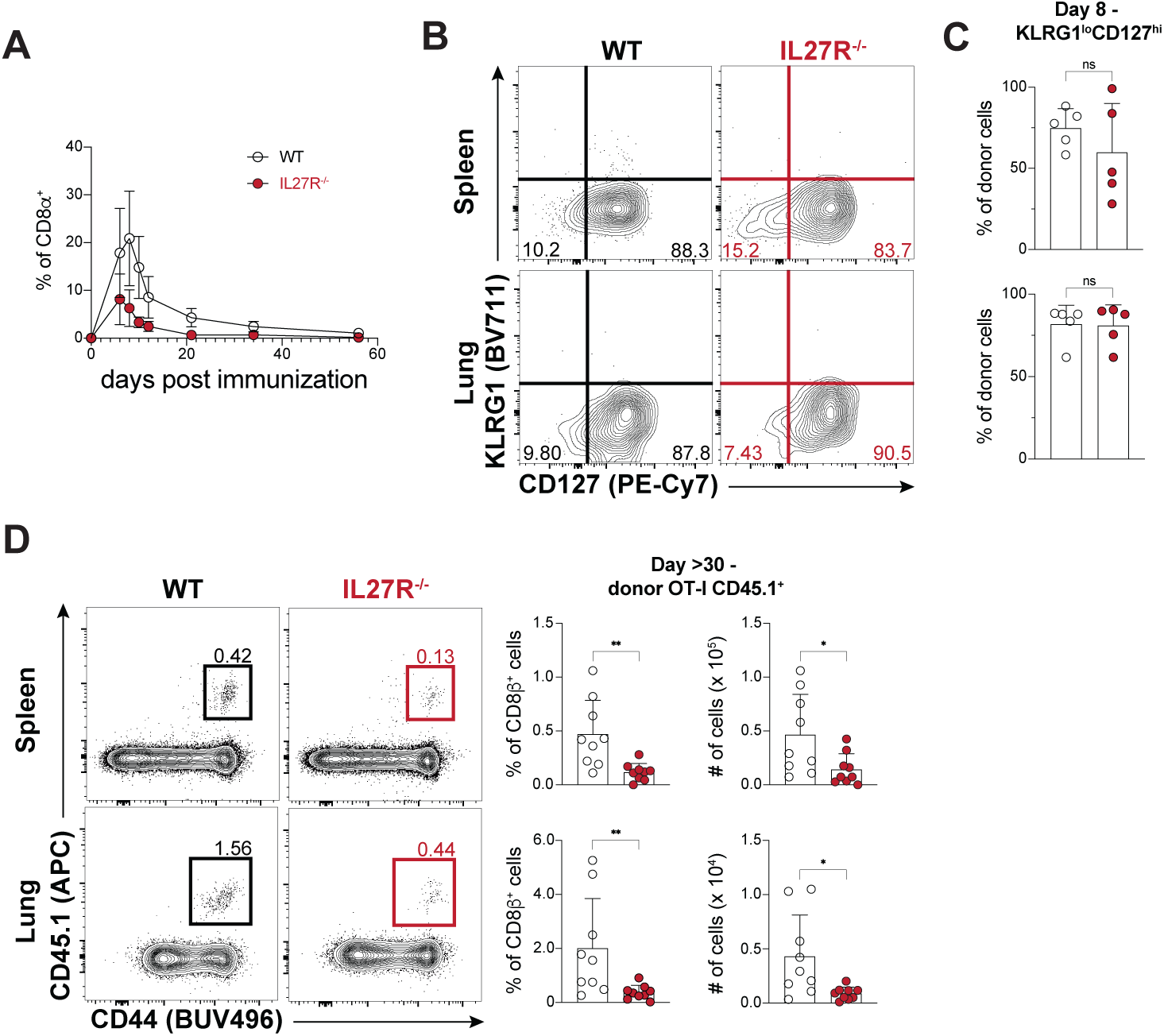
Supporting data that direct IL-27 signaling is necessary for mRNA-LNP induced CD8^+^ T cell responses. **A** Representative analysis of the proportion of donor WT and IL27R^-/-^ OT-Is observed in peripheral blood of WT host mice at indicated days post-immunization. **B-C** Representative flow cytometry analysis of surface phenotype of donor WT and IL27R^-/-^ OT-Is by expression of KLRG1 and CD127 8 days post-immunization in spleen (top) and lungs (bottom). **B** Representative flow cytometry plots of surface KLRG1 and CD127 expression. **C** Summary graphs of proportion of KLRG1^lo^CD127^hi^ subset of donor WT and IL27R^-/-^ OT-Is in spleen (top) and lungs (bottom). **D** Flow cytometry analysis of donor WT and IL27R^-/-^ OT-Is in spleen (top) and lungs (bottom) of WT host mice ≥ 30 days post-immunization. Representative flow cytometry plots of congenically distinct donor OT-Is (left) and summary graphs of proportion and absolute number of indicated donor OT-Is (left). Representative data shown of three experiments of n = 5 mice for each experimental group **A-C.** Cumulative summary data from two experiments of n = 4-5 mice for each experimental group **D**. Data are mean ± SD. ns p > 0.05, * p < 0.05, ** p < 0.01 following Mann-Whitney non-parametric t-test.

**Extended Data Figure 3.**
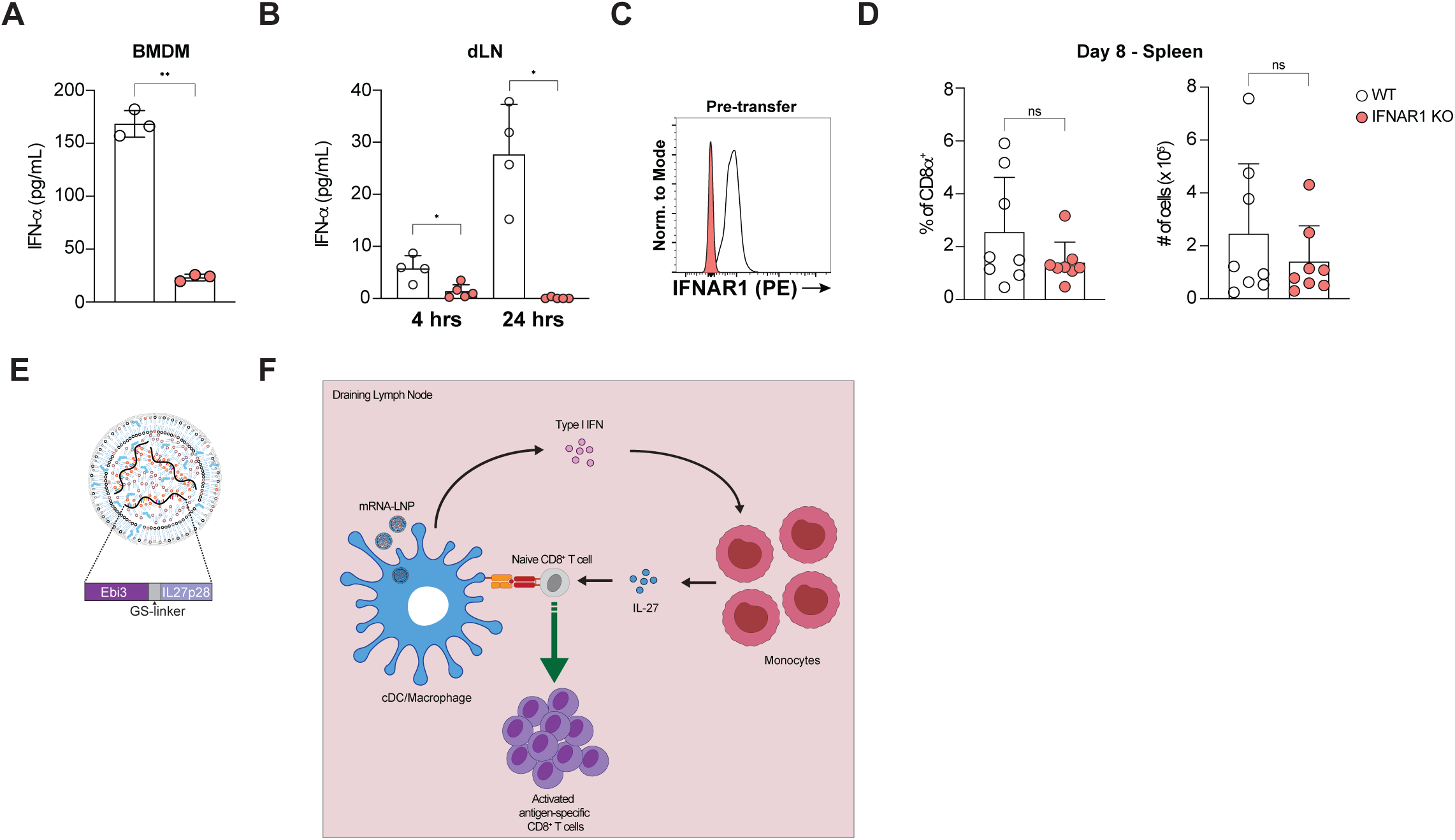
Supporting data that Type I IFN signaling promotes mRNA-LNP induced CD8^+^ T cell responses via IL-27. **A** Production of IFN-α measured by ELISA of WT (open circles) and IFNAR1^-/-^ (filled orange) BMDM lysates and supernatants after overnight incubation with mRNA-LNPs. **B** Production of IFN-α measured by ELISA of WT (open circles) and IFNAR1^-/-^ (filled orange) draining LN lysates 4 hrs and 24 hrs post-immunization with 1 μg of LNP-OVA. **C** Flow cytometry confirmation of IFNAR1 expression on WT and IFNAR1^-/-^ OT-Is prior to adoptive transfer into congenically distinct WT host mice. **D** Analysis of donor WT and IFNAR1^-/-^OT-Is 8 days post-immunization following adoptive transfer into wildtype host mice. Summary graphs of proportion and absolute number of WT and IFNAR1^-/-^ OT-Is identified in the spleen of host mice 8 days post-immunization. **E** Schematic of mRNA-LNPs formulated with IL-27 mRNA consisting of a single mRNA encoding Ebi3 and IL27p28 linked by a short glycine-serine linker. **F** Model schematic of Type I IFN -> IL-27 signaling loop that promotes signaling to CD8^+^ T cells and facilitates expansion following mRNA-LNP immunization. **A-C** Representative data of two independent experiments. n = 4-5 mice per experimental group for **B,D**. **D** Cumulative summary data of two independent experiments. Data are mean ± SD. ns p > 0.05, * p < 0.05, ** p < 0.01. Unpaired Welch’s t-test performed for **A**. Mann Whitney nonparametric t-test performed for **B** and **D**.

**Extended Data Figure 4.**
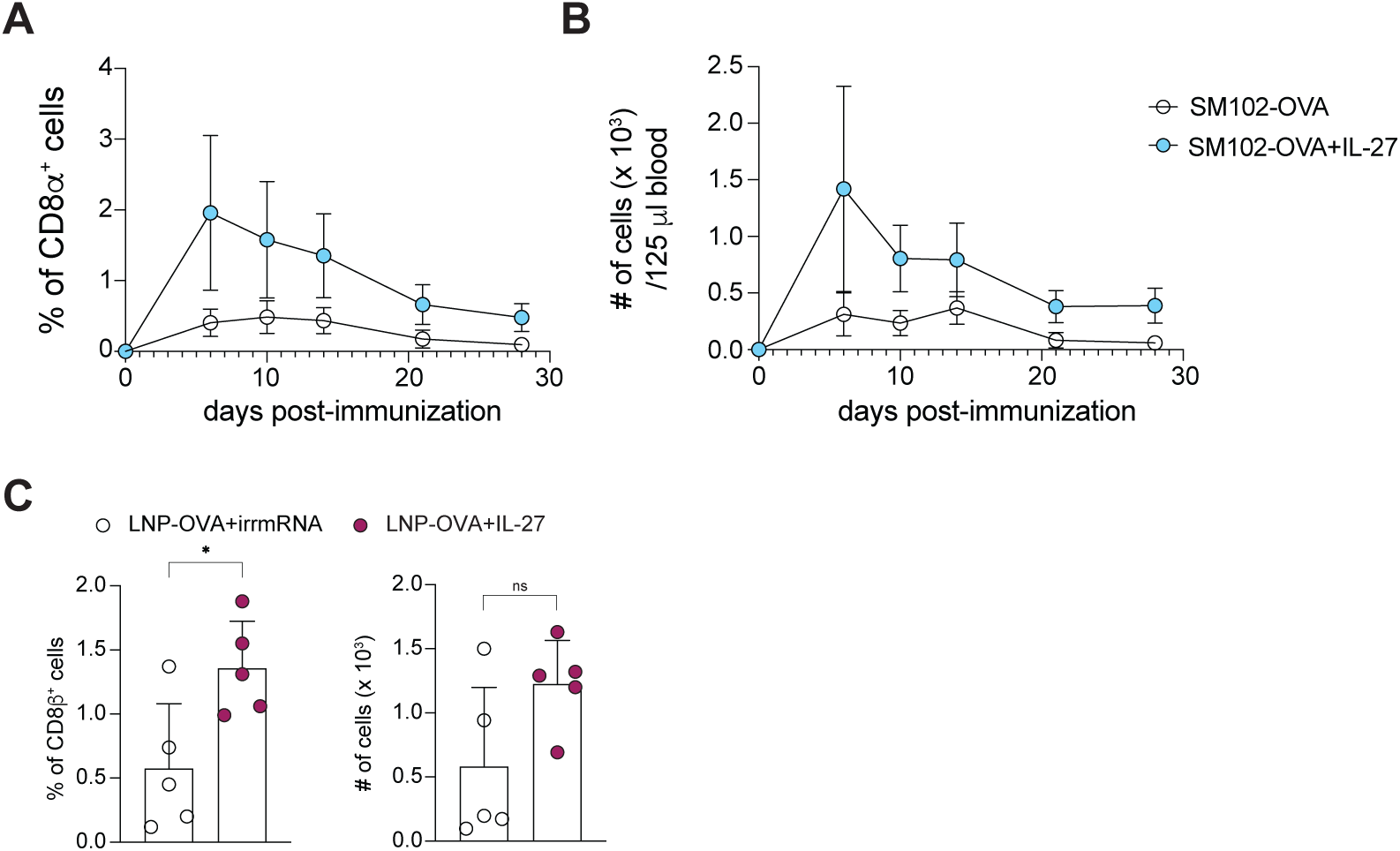
Supporting data that IL-27 mRNA improves the protective capacity of mRNA-LNP elicited CD8^+^ T cells. **A-B** Kinetic analysis of WT OT-Is in peripheral blood of mice at indicated time points following immunization with 1 μg of SM102-OVA or SM102-OVA+IL-27. **A** Donor OT-Is as a proportion of host CD8^+^ T cells. **B** Absolute number of donor OT-I CD8^+^ T cells at each indicated time point. **C** Representative analysis of proportion and absolute number of H2-K^b^-SIINFEKL tetramer^+^ CD8^+^ T cells in peripheral blood of WT mice immunized with LNP-OVA (open) or LNP-OVA+IL-27 (maroon filled) at least 28 days post-immunization and prior to *in vivo* CTL assay. Representative data from two independent experiments of n = 5 mice per experimental group. Data are mean ± SD. ns p > 0.05, and * p < 0.05. Mann Whitney nonparametric t-test performed for **C**.

## Notes

### Competing Interest Statement

In accordance with the University of Pennsylvania and Childrens Hospital of Philadelphia policies and procedures and our ethical obligations as researchers, we report that A.T.P, E.A., M-G.A., D.W., and C.A.H. have filed a provisional patent application that describes the use of nucleoside-modified mRNA encoding cytokines as adjuvants. D.W. is named on patents that describe the use of nucleoside-modified mRNA as a platform to deliver therapeutic proteins and vaccines. D.W. and M-G.A. are named on patents describing the use of lipid nanoparticles, and lipid compositions for nucleic acid delivery and vaccination. We have disclosed those interests fully to the University of Pennsylvania and the Children Hospital of Philadelphia, and we have in place an approved plan for managing any potential conflicts arising from licensing of our patents. Y.T. is an employee of Acuitas Therapeutics and is named on patents describing the use of lipid nanoparticles for nucleic acid delivery. M-G.A. serves as a scientific adviser for AfriGen Biologics and has an ownership stake in RNA Technologies. D.W. serves as a scientific advisor for Arcturus Therapeutics, Cabaletta Bio, and Versatope Therapeutics, and has ownership stakes in Capstan Therapeutics, Orbital Therapeutics, Zipcode Bio, and RNA Technologies. D.W. receives royalties from CellScript and Capstan Therapeutics. R.H.V. has received consulting fees from BMS and Grey Wolf Therapeutics, research funding from Revolution Medicines, is an inventor on patents relating to cancer cellular immunotherapy, cancer vaccines, and KRAS immune epitopes, and receives royalties from Childrens Hospital Boston for a licensed research-only monoclonal antibody. The remaining authors declare no competing interests.

